# TRIM9 Determines Sex-Specific β-Amyloid/Cellular Prion Protein/mGluR5 Complex Formation and Pathological Signaling in Alzheimer’s Disease Mice

**DOI:** 10.64898/2026.07.30.741783

**Authors:** Fatemeh Babaei, Fatemeh A. Panahi, Tash-Lynn L. Colson, Hang Cheng, Mario Tiberi, Rony Chidiac, Stephane Anger, Stephanie L. Gupton, Khaled S. Abd-Elrahman, Stephen S. G. Ferguson

## Abstract

Biological sex is a major determinant of Alzheimer’s disease prevalence, yet the molecular mechanisms underlying sex-specific vulnerability remain largely unknown. Metabotropic glutamate receptor 5 (mGluR5) functions as a co-receptor for β-amyloid (Aβ42) oligomer/cellular prion protein (PrP^C^)-mediated pathogenic signaling in males but not females, establishing a sex-dimorphic node in β-amyloid pathophysiology whose regulatory basis is undefined. Using quantitative proteomic analysis, we identify the E3 ubiquitin ligase TRIM9 as a novel mGluR5-interacting protein and a previously unrecognized sex-specific regulator of the Aβ42/PrP^C^/mGluR5 complex. TRIM9 selectively associates with mGluR5 in male but not female APP/PS1 mouse brain and is required for mGluR5 to serve as a co-receptor for PrP^C^-dependent Aβ42 oligomer binding. Genetic deletion of TRIM9 abolishes Aβ42/PrP^C^/mGluR5 complex assembly in male APP/PS1 mice demonstrating that TRIM9 is an essential scaffold for male-specific Aβ42 signal transduction. Loss of TRIM9 in males further reduces β-amyloid pathology by restoring Akt/GSK3β/ZBTB16-dependent autophagic flux, linking disruption of this complex to a defined downstream proteostatic mechanism. Together, these findings establish TRIM9 as a critical molecular determinant coupling male-specific Aβ42/PrP^C^/mGluR5 complex assembly to downstream neurodegenerative signaling and β-amyloid pathology. They reveal an unappreciated layer of sex-dependent complexity in mGluR5 pharmacology and identify disruption of the TRIM9/mGluR5 interaction as a potential male-specific therapeutic strategy for Alzheimer’s disease.

**Significance Statement:** The molecular basis of sex differences in Alzheimer’s disease vulnerability remains unresolved. We show that mGluR5 functions as a male-specific co-receptor for pathogenic Aβ42/PrP^C^ signaling and identify the E3 ubiquitin ligase TRIM9 as the factor governing this dimorphism. TRIM9 selectively assembles the Aβ42/PrP^C^/mGluR5 complex in male brain, and its genetic deletion disrupts complex formation while restoring Akt/GSK3β-dependent autophagic clearance of amyloid. These findings define a sex-specific signaling axis underlying β-amyloid pathogenesis and establish TRIM9 as a candidate target for sex-informed Alzheimer’s therapeutics.

## Introduction

Metabotropic glutamate receptor 5 (mGluR5) is a key pathological mediator in multiple neurodegenerative diseases, and selective mGluR5 allosteric modulators can either reverse or slow memory and motor deficits in mouse models of amyotrophic lateral sclerosis, Parkinson’s disease, Huntington’s disease, and Alzheimer’s disease (AD)^1–12^. In AD specifically, synaptic dysfunction, memory loss and cognitive decline track closely with the progressive accumulation of two neurotoxic species: soluble β-amyloid (Aβ42) oligomers and hyper-phosphorylated tau^13–15^. Critically, Aβ42 oligomers disrupt synaptic plasticity and memory in AD prior to significant neuronal loss through their interaction with the high-affinity GPI-anchored receptor cellular prion protein (PrP^C^) and by promoting the clustering and pathological activation of mGluR5-mediated Ca^2+^ release^16–21^. Specifically, mGluR5 operates as an essential co-receptor protein that couples Aβ42/PrP^C^ to pathological intracellular signaling cascades that underlie AD pathology^1,3,4,6,22^.

mGluR5 allosteric modulators have been the focus of extensive drug discovery efforts and represent promising novel pharmacotherapeutics but have repeatedly faltered in clinical trials because sex as a biological variable is often neglected in preclinical research^12,15,23–25^. This oversight carries particular weight given a striking and counterintuitive discovery that despite the fact that AD disproportionately impacts women, pharmacological blockade of mGluR5 selectively prevents the progression of memory impairment and disease pathology in male but not female APPswe/PS1ΔE9 (APP/PS1) AD mice^8,26^. This male-restricted efficacy reflects a sexual dimorphism that extends throughout the Aβ42/PrP^C^/mGluR5 signaling axis. Aβ42 oligomers bind mGluR5 with high affinity in male, but not female, mouse, rat, and human brain tissue; PrPC forms a physical complex with mGluR5 exclusively in male brain tissue; and mGluR5 antagonism reactivates a GSK3β/ZBTB16/ATG14 autophagy pathway that reduces Aβ pathology only in male APP/PS1 mice⁸,²⁷. Collectively, these findings indicate that the Aβ42/PrP^C^/mGluR5 signaling axis is either absent or functionally uncoupled in females, conferring resistance to both mGluR5-dependent pathology and mGluR5-targeted pharmacological intervention. Notably, although estrogen receptor (ER) antagonism restores cell-surface mGluR5 expression in female APP/PS1 mice but does not rescue GSK3β-dependent autophagic signaling. This indicates that sex differences in this pathway are not attributable to receptor trafficking alone and instead reflect more fundamental differences in the mGluR5 interactome and downstream signaling pathways^28^.

Here, using quantitative proteomics, we identify TRIM9, a brain-specific E3 ubiquitin ligase that is enriched in post-synaptic densities and known to govern hippocampal neuron development, as a novel male-specific mGluR5 regulatory protein in APP/PS1 mouse cortex^29–32^. Genetic deletion of *Trim9* disrupts assembly of the Aβ42/PrP^C^/mGluR5 complex formation and prevents Aβ42/PrP^C^-driven suppression of autophagic flux culminating in reduced Aβ burden in male APP/PS1 mice. Our findings identify TRIM9 is a critical intracellular component of male-specific pathological Aβ42/PrP^C^/mGluR5 signaling uncovering a previously unrecognized mechanism that underlies sex-dependent AD pathology and opening a compelling new avenue for precision, sex-informed therapeutics.

## Results

### Identification of TRIM9 as a component of Aβ42/PrP^C^/mGluR5 signaling complex

We previously reported that Aβ42 oligomers bound solely to mGluR5 in male but not female mouse, rat, and human cortical tissue^8^. Therefore, we performed liquid chromatography-tandem mass spectrometry (LC-MS/MS) analysis to identify proteins that were selectively co-immunoprecipitated with mGluR5 from cortical tissue from either 6-month-old male and female mice. Cortical tissue from male and female mGluR5 knockout mice were employed as controls. A total of 249 proteins were identified across samples. Following removal of known contaminants and filtering based on detection in biological replicates, 36 proteins were retained for downstream analysis. Proteins identified in mGluR5 immunocomplexes included components associated with synaptic signaling and membrane trafficking, including RasGRF2, clathrin heavy chain, Homer1 and TRIM9 (Fig. 1a). Selective interactions with mGluR5 in male versus female cortical tissue was visualized using a volcano plot with only TRIM9 and Hook3 proteins demonstrating statistically significant selectivity for mGluR5 in male cortex with an absolute log₂ fold change ≥ 1 and FDR < 0.05 (Fig. 1b).

**Fig. 1.**
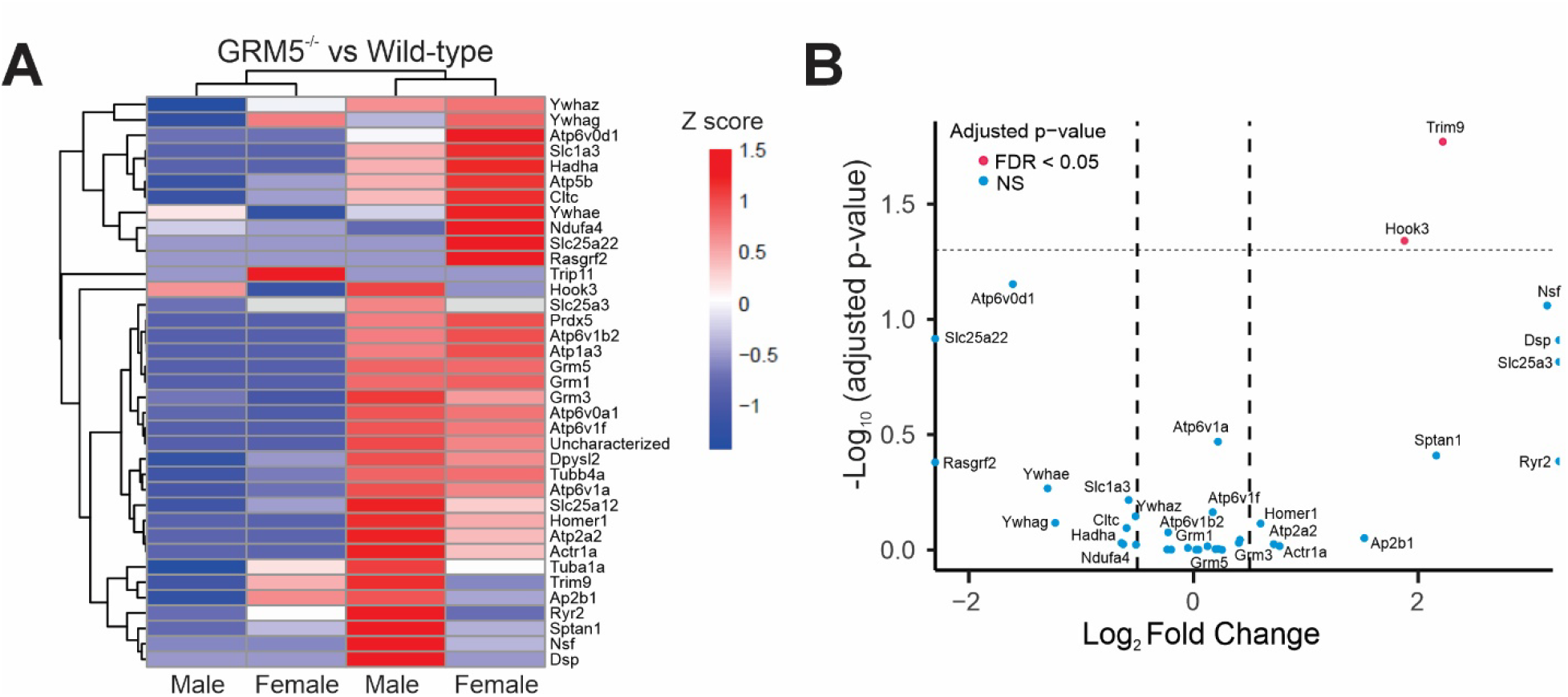
Proteomic identification of TRIM9 as a male specific mGluR5 interacting protein. **(a)** LFQ protein intensity were averaged across biological replicates per condition, and row-wise Z-score normalization was applied to visualize relative abundance patterns across samples. Z-scored values were used exclusively for visualization and clustering. Data were grouped as the mean of values for each experimental condition (6-month-old wild-type males n=4, 6-month-old GRM5^-/-^ males n=3, 6-month-old wild-type females n=3, 6-month-old GRM5^-/-^ females n=3). **(b)** LFQ intensity values were log2-transformed and processed using Perseus, including missing value imputation. Differential protein abundance was assessed using one-way ANOVA followed by Tukey’s post-hoc test, with multiple testing correction applied to p-values. The volcano plot shows log2 fold change (x-axis) versus –log10 p-value (y-axis) for the 6-month-old male versus female comparison. Significance thresholds are indicated by dashed lines, and significantly altered proteins (p< 0.05) are highlighted.

TRIM9 was previously identified as a brain-specific E3 ubiquitin ligase that was enriched in post-synaptic densities and played an important role in neuronal development in the hippocampus and cortex^29–32^. The association of TRIM9 with mGluR5 was confirmed by co-immunoprecipitation of mGluR5 from 6-month-old male and female wild-type and APP/PS1 cortex using male and female *GRM5^-/-^* and *Trim9^-/-^* mouse cortex as negative controls. We found that both the long (∼80kDa) and short (∼60 kDa) isoforms of TRIM9 were co-immunoprecipitated with mGluR5 from wild-type male and female cortex (Fig. 2a). In contrast, TRIM9 co-immunoprecipitated with mGluR5 from male APP/PS1 cortex but the interaction between mGluR5 and TRIM9 was lost in female APP/PS1 mouse cortex (Fig. 2a). Similarly, mGluR5 was co-immunoprecipitated with TRIM9 from both wild-type male and female cortex but was only co-immunoprecipitated with TRIM9 from male APP/PS1 mouse cortex (Fig. 2b). We assessed which intracellular mGluR5 domains were contributing to TRIM9 binding to mGluR5 in transfected HEK293T cells and found that the substitution of lysine residues at positions 690–692 within the intracellular loop 2 (IL2) region of mGluR5 with arginine residues (K690R, K691R, K692R) reduced the association between TRIM9 and mGluR5 (Fig. 2c). We previously demonstrated that the identical residues in mGluR1 contributed to G protein-coupled receptor kinase 2 (GRK2) association with the receptor^33^. When assessed by co-immunoprecipitation in transfected HEK293T cells, we found that the TRIM9 interaction with mGluR5 was reduced following the overexpression of GRK2 (Fig. 2d). Taken together these findings indicated that TRIM9 forms a complex with mGluR5 via its association with IL2 and that the interaction was male-selective under pathological conditions in APP/PS1 mice as interactions between TRIM9 and mGluR5 were lost in female APP/PS1 mice.

**Fig. 2.**
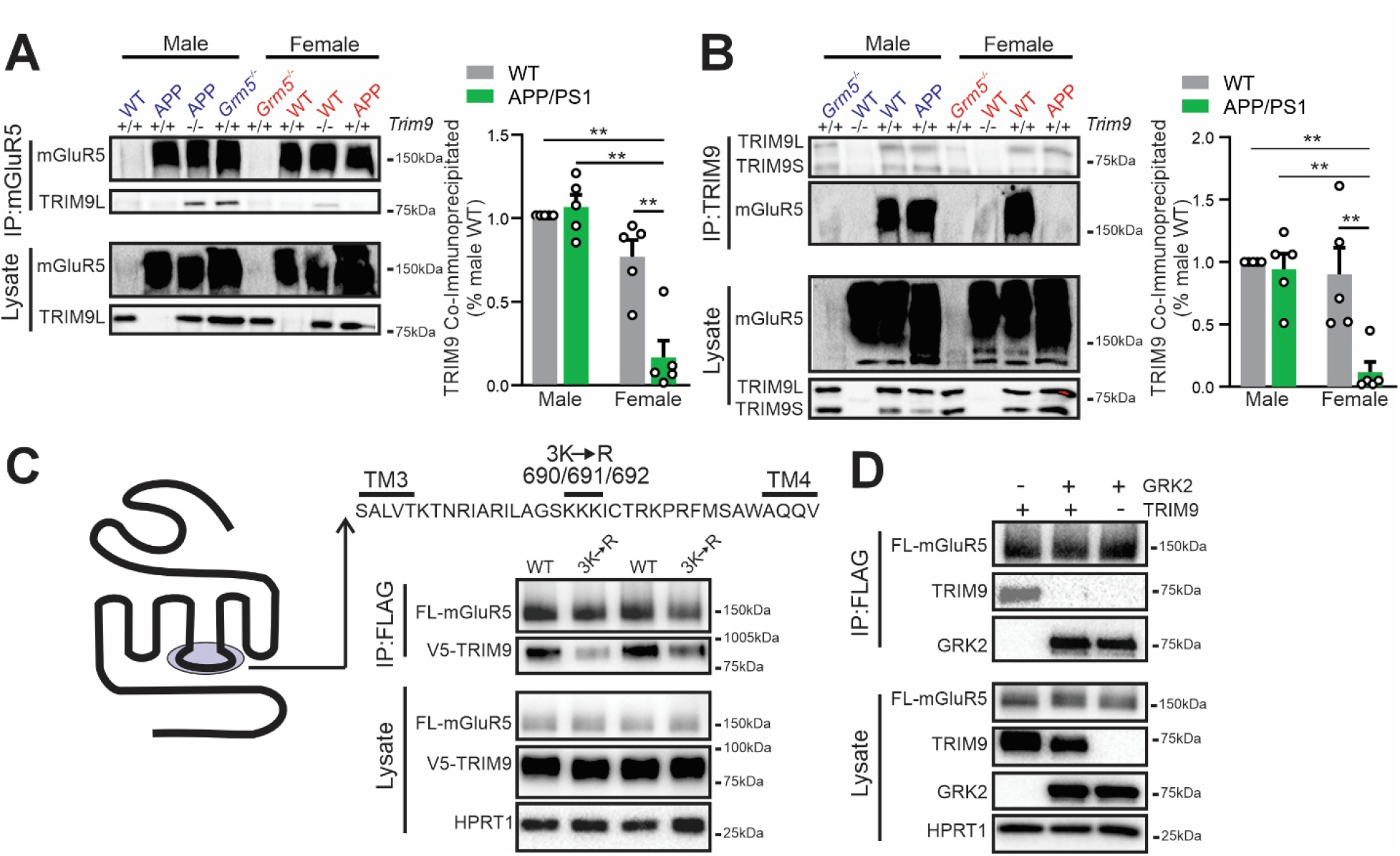
TRIM9 is a component of Aβ42/PrP^C^/mGluR5 signaling complex in 6-month-old male and female mice. **(a)** Representative immunoblots and quantitative analysis of the co-immunoprecipitation of TRIM9 with mGluR5 from cortical tissue (500 µg) from 6-month-old male and female wild-type (WT), *Trim9^-/-^*, APP/PS1 and APP/PS1::*Trim9^-/-^* mice. Data represent the mean ± SEM (n = 5 mice per group). Statistical analysis was assessed using one-way ANOVA (F(3,16)=26.93, P<0.0001) followed by Tukey’s multiple comparisons test. **p<0.01. **(b)** Representative immunoblots and quantitative analysis of the co-immunoprecipitation of mGluR5 with TRIM9 from cortical tissue (500 µg) from 6-month-old male and female WT, *Trim9^-/-^*, APP/PS1 and APP/PS1::*Trim9^-/-^* mice. Data represent the mean ± SEM (n = 5 mice per group). Statistical analysis was assessed using one-way ANOVA (F(3,16)=10.25, P=0.0005) followed by Tukey’s multiple comparisons test. **p<01. **(c)** Representative replicates V5-tagged TRIM9 co-immunoprecipitation with either FLAG-mGluR5 or FLAG-mGluR5-K690R/K691R/K692R in HEK293T cells. (n=4). **(d)** Representative co-immunoprecipitation of V5-TRIM9 with FLAG-mGluR5 in the presence and absence of GRK2 from transfected HEK293T cells (n=3).

### TRIM9 deletion disrupts the male-specific Aβ42/PrP^C^/mGluR5 protein complex

We previously found that while Aβ42 oligomers bind to mGluR5 in male mouse cortex and hippocampus with sub-nanomolar affinity, whereas Aβ42 oligomers exhibited no affinity for the receptor in female brain tissue^8^. Therefore, we examined whether Aβ42 oligomer binding to mGluR5 was altered in cortical membranes from six-month-old male and female *Trim9^-/-^* mice using [^3^H]-MPEP radioligand binding assays. Aβ42 oligomers displaced [^3^H]-MPEP binding in cortical membranes from wild-type male mice, but not from *Trim9^-/-^*male mice (Fig. 3a). Aβ42 oligomers did not displace [^3^H]-MPEP binding to either wild-type or *Trim9^-/-^* female mouse cortical membranes, which is in agreement with our previous studies (Fig. 3b)^8^. Aβ42 oligomers were previously shown to promote the interaction of PrP^C^ with mGluR5 in mouse brain preparations from APP/PS1 mice and PrP^C^ was not observed to co-immunoprecipitate with mGluR5 from female mouse hippocampus^4,34^. Therefore, we examined whether *Trim9^-/-^* deletion altered PrP^C^ interactions with mGluR5 in 6-month-old male and female wild-type and APP/PS1 mouse cortex. PrP^C^ did not co-immunoprecipitate with mGluR5 from either female wild-type or APP/PS1 cortex in either absence of presence of TRIM9 expression (Fig. 3c and 3d). In agreement with previous work^4,34^, we found that PrP^C^ interactions were significantly enhanced in male APP/PS1 cortex when compared to control mice (Fig. 3c and d). However, while *Trim9* deletion did not prevent PrP^C^ co-immunoprecipitation with mGluR5 from the cortex of wild-type male mice, the increased mGluR5/PrP^C^ interactions observed in male APP/PS1 mouse brain were reduced back to wild-type levels in the absence of TRIM9 expression (Fig. 3c and d). These observations suggest that TRIM9 contributes directly to the formation of Aβ42/PrP^C^/mGluR5 signaling complexes (Fig. 3e).

**Fig. 3.**
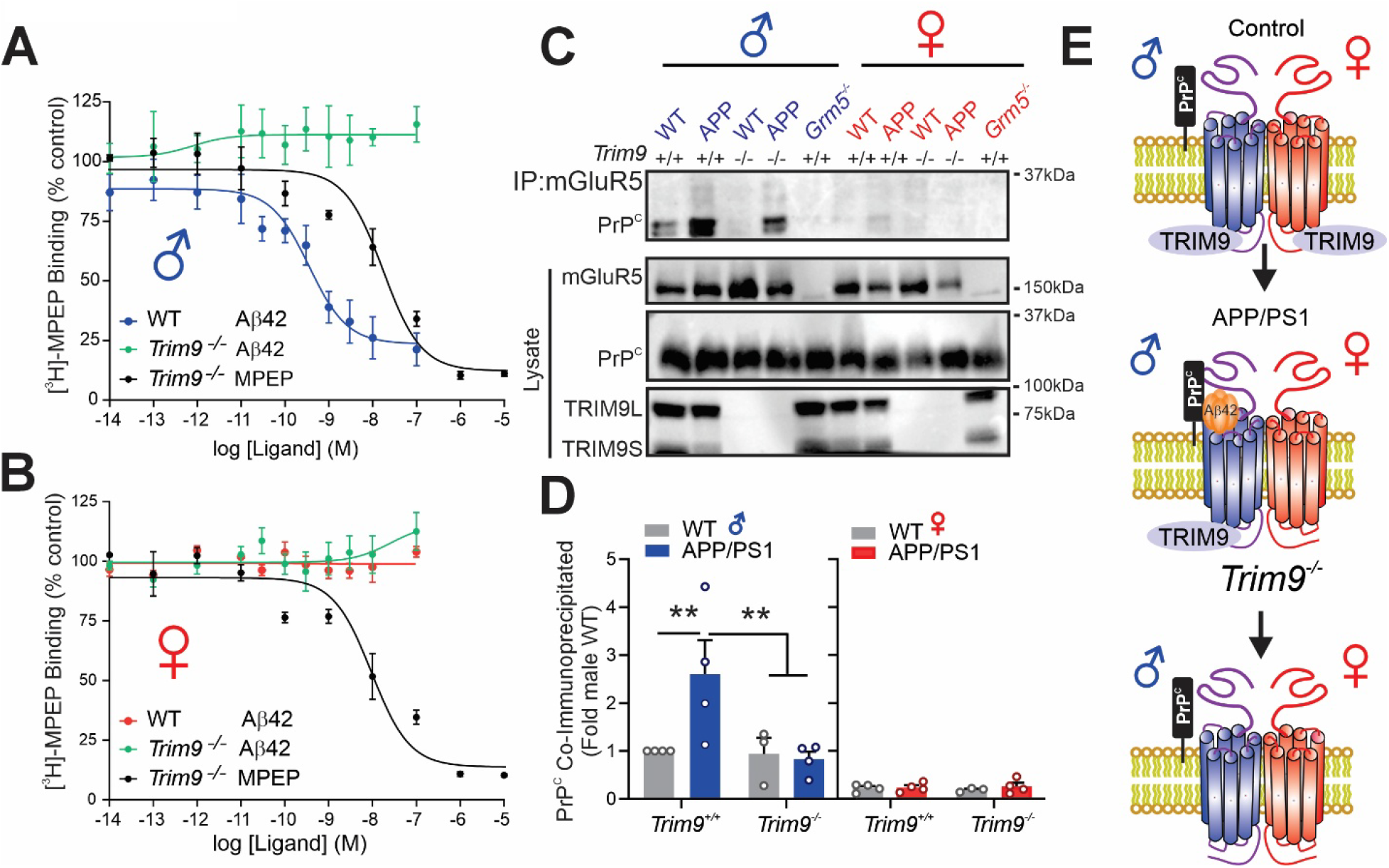
*Trim9* deletion disrupts the male-specific Aβ42/PrP^C^/mGluR5 protein complex a formation. [³H]-MPEP displacement curves for MPEP and Aβ42 for WT and *Trim9^-/-^* male **(a)** and female **(b)** mouse cortex. Each binding curve represents the mean ± SEM of technical duplicates (n=3-4). Representative immunoblots **(c)** and quantitative analysis **(d)** of PrP^C^ co-immunoprecipitation with mGluR5 from cortical tissue (500 µg) from male and female WT, *Trim9^-/-^*, APP/PS1 and APP/PS1::*Trim9^-/-^* mice. Data are presented as the mean ± SEM (n=3-4 mice per group). Statistical analysis was assessed using two-way ANOVA (F(3,22)=4.4, P=0.0145) followed by Tukey’s multiple comparisons test **p<0.01.. **(e)** Schematic representation illustrating sex-specific formation of the Aβ42/PrP^C^/mGluR5 complex and its disruption in the absence of TRIM9 protein expression.

Because pharmacological blockade of mGluR5 resulted in reduced Aβ pathology in male but not female APP/PS1 mice^8^, we tested whether *Trim9* deletion would reduce Aβ pathology in male APP/PS1 mice at 10 months of age. We found that β-amyloid plaque deposition in the hippocampus of male APP/PS1::*Trim9^-/-^* mice was significantly reduced when compared to male APP/PS1 mice (Fig. 4a and b). To further examine whether TRIM9 deletion influences Aβ oligomer levels, Aβ oligomers were quantified in cortical tissue from male wild-type, *Trim9^-/-^*, APP/PS1 and APP/PS1::*Trim9^-/-^* mice. Wild-type APP/PS1 animals displayed markedly elevated Aβ oligomer levels relative to wild-type controls, whereas Aβ oligomer levels in male APP/PS1::*Trim9^-/-^* mice were identical to those found in either wild-type or *Trim9^-/-^*male mice at 10 months of age (Fig. 4c).

**Fig. 4.**
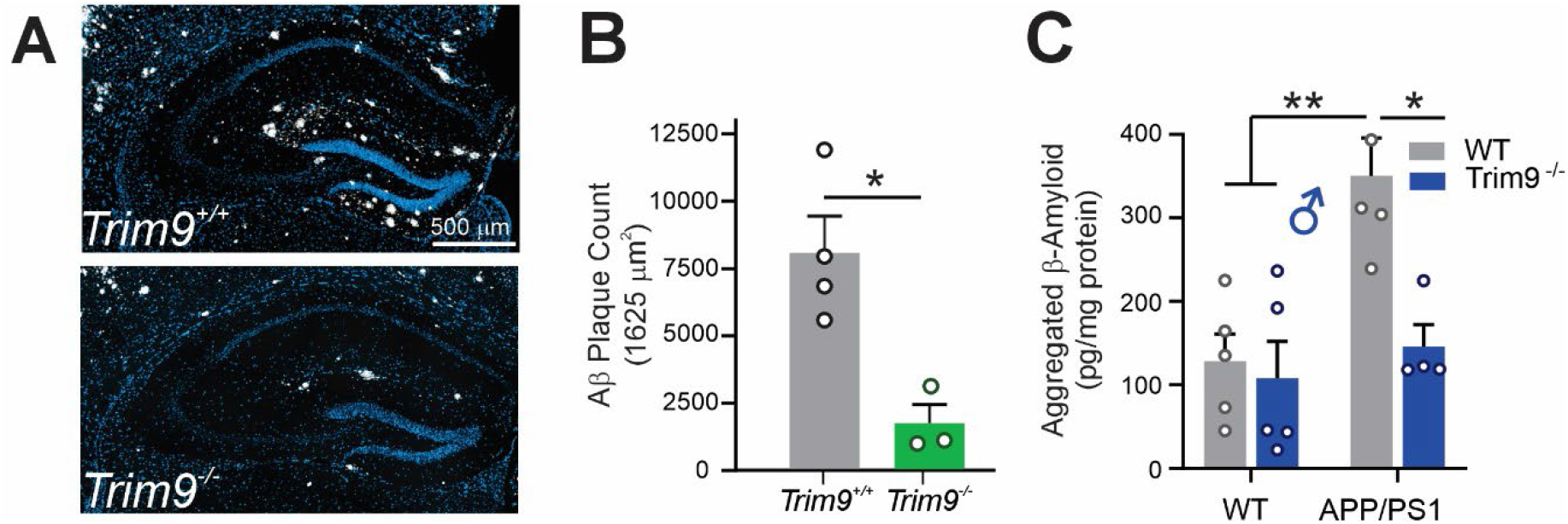
*Trim9* deletion attenuates Aβ pathology in male APP/PS1 mice. **(a)** Representative images of Aâ plaques in APP/PS1 hippocampus in the presence (*Trim9^+/+^*) or absence (*Trim9^-/-^*) expression. **(b)** Quantification of plaque density in 1625 µm^2^ regions of interest in the hippocampus of six brain slices per male APP/PS1 mice in the presence (*Trim9^+/+^*) or absence (*Trim9^-/-^*) expression. Data are presented as mean ± SEM (n =3-4 mice per group). Statistical analysis was assessed by two tailed unpaired T-test. *<0.05 versus APP/PS1 *Trim9^+/+^* mice. **(c)** Quantitative analysis aggregated β-amyloid concentrations (pg/mg protein) in cortical tissue from male WT, *T*rim9^-/-^, and APP/PS1 mice as well as male APP/PS1 mice lacking TRIM9 expression at 10 months of age. Data are presented as mean ± SEM (n = 4-5) mice per group). Statistical analysis was assessed using one-way ANOVA followed by Tukey’s multiple comparisons test (F(3,15)=8.61, P=0.0015). *p<0.05, **p<0.01 as indicated by bars in the Figure.

### TRIM9 associates with GSK3β and regulates downstream kinase signaling to regulate autophagy

TRIM9 was previously reported to function as a scaffold capable of recruiting GSK3β to TBK1 signaling complexes^35^. In both male and female mice, GSK3β was detected in TRIM9 immunoprecipitates from wild-type cortex but not in TRIM9 knockout samples, and remained detectable in *Grm5^⁻/⁻^* tissue, indicating that this interaction does not require mGluR5. (Fig. 5a). GSK3β activity is regulated by Akt-mediated phosphorylation of serine residue 9 and pS9-GSK3β phosphorylation was previously shown to be elevated in male but not female APP/PS1 mice^8^. The phosphorylation of Akt (pS473) and pS9-GSK3β (pS9) is elevated specifically in male APP/PS1 mice and can be blocked by either mGluR5 knockout or by treating male APP/PS1 mice with a mGluR5-specific antagonist^8,34^. Therefore, because TRIM9 interacts with both mGluR5 and GSK3β, we examined whether TRIM9 contributed to the regulation of both pS473-Akt and pS9-GSK3β phosphorylation in male but not female APP/PS1 mice. We found that both pS473-Akt and pS9-GSK3β phosphorylation were selectively increased in male APP/PS1 mouse cortex when compared to either wild-type or *Trim9^-/-^*male mice (Fig. 5b-d). However, the increased phosphorylation of pS473-Akt and pS9-GSK3β observed in male APP/PS1 mice was reduced to wild-type levels in the absence of TRIM9 (Fig. 5c and d). pS473-Akt and pS9-GSK3β phosphorylation levels remained unchanged in female APP/PS1 mice (Fig. 5b-d).

**Fig. 5.**
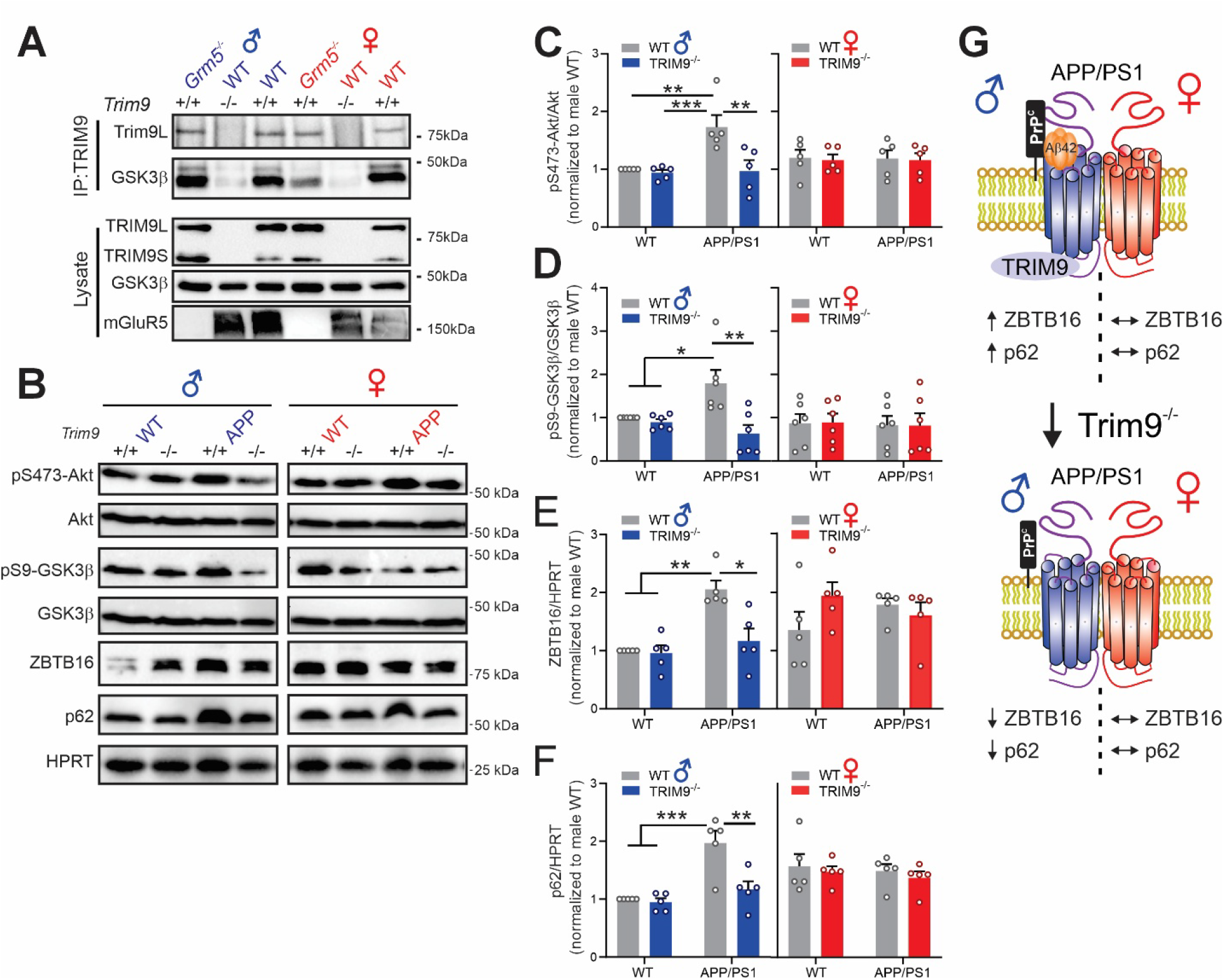
TRIM9 associates with GSK3β and regulates downstream kinase signaling and autophagy. **(a)** Representative immunoblots of GSK3β co-immunoprecipitation with TRIM9 from cortical tissue (500 µg) from 6-month-old male and female WT and *Grm5^-/-^*mice in the presence (+/+) or absence (-/-) of TRIM9. (n=3). Representative immunoblots **(b)** and quantitative analysis of **(c)** pS473-Akt phosphorylation (F(3,32)=4.3, P=0.0114) (n=5), **(d)** pS9-GSK3β phosphorylation (F(3,40)=2.9, P=0.0428) (n=5-6), **(e)** ZBTB16 protein expression (F(3,32)=5.262, P=0.0046) (n=5) and **(f)** p62 protein expression (F(3,32)=5.056, P=0.0022) (n=5) in lysates from cortical tissue (500 µg) from 6-month-old male and female WT, *T*rim9^-/-^, APP/PS1 and APP/PS1::*Trim9^-/-^*mice. Statistical analysis was assessed using two-way ANOVA followed by Tukey’s multiple comparisons test. *<0.05, **p<0.01; ***p<0.001. **(g)** Schematic representation illustrating sex-specific regulation of Akt, GSK3β and ZBTB16 signaling downstream of the mGluR5 in male and female APP/PS1 mice by TRIM9 resulting in reduced p62 expression and Aβ pathology in only male APP/PS1 mice.

Previous studies have also shown that pS9-GSK3β phosphorylation antagonizes a ZBTB16-mediated autophagy pathway in both APP/PS1 and 3xTg AD mice^8,27^. Therefore, we examined whether TRIM9 contributed to the regulation of pS9-GSK3β phosphorylation-dependent regulation of ZBTB16 protein expression in male and female APP/PS1 mice. Consistent with previous findings in 12-month-old APP/PS1 mice^8^, we found that ZBTB16 protein expression was significantly increased in the cortex of 6-month-old male but not female APP/PS1 mice (Fig. 5b and e). However, ZBTB16 protein expression was reduced back to control levels in male APP/PS1 mice lacking TRIM9 expression (Fig. 3A and E). Moreover, p62 levels were increased only in male APP/PS1 mice and were reduced to wild-type levels in male APP/PS1 mice lacking TRIM9 expression (Fig. 5b and f). Thus, TRIM9 also contributes to male-specific mGluR5-mediated inactivation of GSK3β signaling in APP/PS1 mice and we find that reductions in Aβ pathology are paralleled by the reactivation of a GSK3β/ZBTB16 autophagic pathway in male APP/PS1 mice (Fig. 5g)^8,36^.

## Discussion

We show here that TRIM9 is a pivotal and previously unrecognized regulator of pathological Aβ42 signaling in AD that functions to both promote extracellular Aβ42/PrP^C^/mGluR5 complex formation and transduce receptor-mediated intracellular pathological signals. In male APP/PS1 mice, genetic deletion of *Trim9* abolishes the ability of both Aβ42 to associate with mGluR5 in both wild-type and APP/PS1 male mouse cortex. PrP^C^ interactions with mGluR5 are enhanced in male APP/PS1 mice when compared with control mice, but the increase in PrP^C^/mGluR5 complex formation is lost in male APP/PS1 mice lacking TRIM9. Strikingly, neither Aβ42 nor PrP^C^ interact with mGluR5 in female APP/PS1 mice, regardless of TRIM9 expression, highlighting a profoundly male-specific mechanism underlying pathological signaling in AD. Beyond regulating receptor complex formation, TRIM9 also biases mGluR5 signaling toward disease-associated pathways by promoting the inactivation of Akt-, and GSK3β-dependent signaling cascades. These pathological signaling events suppress autophagic flux and ultimately facilitate Aβ accumulation in male AD mice. These findings identify TRIM9 as a central mediator of a male-specific Aβ42/PrP^C^/mGluR5 signaling by linking extracellular mGluR5 signaling to the assembly of downstream intracellular signal transduction pathways underlying neurodegenerative pathology in AD. This provides a mechanistic explanation for how mGluR5 acquires male-specific pathological function in AD.

In cortical and hippocampal tissue from female mice, both Aβ42 and PrP^C^ fail to associate with mGluR5. Consequently, mGluR5-mediated alteration in Akt, and GSK3β signaling that occurs in male APP/PS1 mice that are affected by the genetic deletion of *Trim9*, are not observed in female APP/PS1 mice^2,8,37^. While TRIM9 can be co-immunoprecipitated with mGluR5 from control but not APP/PS1 female cortical tissue, both Aβ42 and PrP^C^ fails to interact with mGluR5 in control female brain tissue. This suggests that an additional component contributes to the protection of female mice from pathological β42/PrP^C^/mGluR5/TRIM9 signaling. Specifically, increased interactions between plasma membrane ERα and mGluR5 have been reported in female PTEN knockout mice resulting in increased mGluR5-dependent seizure activity^38^. Moreover, both pharmacological inhibition ERα and the failure of ERα to express at the cell surface results in a gain of function phenotype for Aβ42/mGluR5-mediated suppression of GIRK channel activity^39^. Whether an ERα-dependent mechanism underlies the sex-specific mGluR5 pathophysiology observed in AD mice remains to be determined, but ERα/mGluR5 interactions likely protect against mGluR5-mediated pathology resulting a reduction in mGluR5/TRIM9 interactions in female AD mice. Regardless, attempts to modify putative interactions of ERα with mGluR5 would not be of therapeutic benefit to female AD patients.

We find that both pS473-Akt and pS9-GSK3β phosphorylation are increased in male but not female APP/PS1 mouse cortex. The increase in Akt and GSK3β phosphorylation results in the inactivation of a ZBTB16-dependent autophagic pathway and the accumulation of misfolded protein (p62) linked to increases in protein aggregates in both AD and Huntington’s disease mice^8,36,37,40–42^. However, both genetic deletion of *Trim9* and pharmacological inhibition of mGluR5 reverse the elevations in Akt and GSK3β phosphorylation that we show are specific to male APP/PS1 mice^8,36,42^. Akt- and GSK3β-regulated autophagy pathways converge as the consequence of Akt-mediated phosphorylation of GSK3β at Ser9 resulting in the accumulation of ZBTB16, a ubiquitin ligase that regulates the expression of the autophagy adaptor protein ATG14^8,36,43^. TRIM9 functions as a GSK3β scaffolding protein and we propose that TRIM9 interactions with the second intracellular loop domain mGluR5 allow it to functionally co-ordinate phosphorylation and inactivation of Akt and GSK3β in response to Aβ42/PrP^C^-mediated activation of mGluR5. Interestingly, Akt and GSK3β form a complex with protein phosphatase 2A and β-arrestin2^44,45^. While no studies have shown a direct interactions between mGluR5 and β-arrestin2, β-arrestin2 can be co-immunoprecipitated with mGluR5 and contributes to downstream mGluR5 signaling and pathology in Fragile X mice^46,47^. Thus, TRIM9 may function as molecular intermediate to organize β-arrestin2-biased signaling downstream of mGluR5.

In this work, we have identified TRIM9 as a novel component of the Aβ42/PrP^C^/mGluR5 signaling axis that selectively enables male-specific engagement of mGluR5 with both Aβ42 and PrP^C^. TRIM9 further functions as an intracellular component of the Aβ42/PrP^C^/mGluR5 complex in male AD mice to couple to the receptor to the pathological dysregulation of autophagy resulting Aβ pathology. Our results suggest that TRIM9 is a essential intracellular component of a male-specific pathological signaling pathway and establishes a molecular framework that explains how identical extracellular AD-associated stimuli can produce fundamentally different biological outcomes between the sexes. These findings highlight the complexity of mGluR5 pharmacology and Aβ signaling and identify TRIM9/mGluR5 interactions as a male specific target for AD therapeutics.

## Materials and Methods

### Animals

APP/PS1::*Trim9^-/-^* mice were generated by crossing B6;C3-Tg(APPswe,PSEN1dE9)85Dbo/Mmjax mice (strain 004462) purchased from Laboratory on a C57BL/6 background with *Trim9^-/-^* knockout mice maintained on the same background (29-32). Offspring were identified by ear notch biopsy and genotyped by polymerase chain reaction (PCR) using primers specific for the APP transgene and the *Trim9* allele^2,29–32^. The following genotypes were obtained and used for experiments: wild-type, APP/PS1, *Trim9^-/-^,* and APP/PS1:: *Trim9^-/-^*. Mice were housed in groups of four per cage under standard conditions with ad libitum access to food and water, maintained on a 12 h light/dark cycle at 24°C. Mice were aged to 6 months, unless otherwise indicated in the *Figure Legends*, and then euthanized by exsanguination. Brains were rapidly collected and processed for biochemical analyses. All animal experiments were reviewed and approved by the University of Ottawa Institutional Animal Care Committee and carried out in accordance with Canadian Council on Animal Care guidelines.

### Antibodies and Reagents

Primary and secondary antibodies used in this study were obtained from multiple commercial sources. Rabbit anti-GSK3β (phospho-Ser9, #9323), anti-Akt (phospho-Ser473, #4060), anti-ULK1 (phospho-Ser757, #14202), anti-mTOR (phospho-Ser2448, #109268), and total mTOR (#2972), as well as mouse anti-Akt (#9272) and anti-GSK3β (#9832) antibodies were obtained from Cell Signaling Technology (Massachusetts, USA). Mouse anti-p62 (#56416), anti-PrPᶜ (ab61409), and rabbit anti-ZBTB16 (#39354) were obtained from Abcam (Cambridgeshire, UK). Rabbit anti-mGluR5 (AB5675) antibody, Anti-FLAG M2 affinity agarose beads and protease inhibitors cocktail (539134) were obtained from Millipore Sigma (Massachusetts, USA). Rabbit anti-HPRT1 (MA5-32797) was purchased from Thermo Fisher Scientific (Massachusetts, USA). Rabbit anti-ULK1 (20986-1-AP) and anti-TRIM9 (10786-1-AP) antibodies were purchased from Proteintech (Illinois, USA). Protein G Mag Sepharose beads (28-9513-79) were purchased from Cytiva (Massachusetts, USA). Western blotting reagents were purchased from Bio-Rad (California, USA). For cell culture experiments, Dulbecco’s Modified Eagle Medium (DMEM) and sterile phosphate-buffered saline (PBS) were obtained from Wisent Bioproducts (Quebec, Canada). Fetal bovine serum (FBS) was purchased from Corning (New York, USA). Penicillin– streptomycin, 0.25% trypsin-EDTA, and Opti-MEM reduced serum medium were obtained from Thermo Fisher Scientific (Massachusetts, USA). Enhanced polyethylenimine (ePEI) used for transfection was obtained from the Genome Editing and Molecular Biology Facility, University of Ottawa. Poly-D-lysine (PDL) and all other biochemical reagents were purchased from Millipore Sigma (Massachusetts, USA).

### LC-MS/MS analysis

Proteins associated with mGluR5 were subjected to enzymatic digestion and analyzed by liquid chromatography–tandem mass spectrometry (LC–MS/MS). For protein digestion, 0.2 µg of Trypsin/Lys C (V5071, Promega) was added into each 100 µl sample in 50 mM ammonium bicarbonate overnight at 37 °C. Supernatants were transferred into new microtubes, beads were washed two times with 100 µl of water and supernatants from all washes were pooled with the initial supernatant. Trifluoroacetic acid (1%; 302031 Sigma) was added into each tube to acidify samples. Samples were then desalted using desalting spin columns (89852; Pierce, Thermo Fisher) per manufacturers protocol and subsequently dried in a SpeedVac. Peptides were resuspended in 26 µl of 1% formic acid and kept at -80°C. Peptides were loaded onto a 75-μm internal diameter × 150 mm Self-Pack C18 column installed in an Easy-nLC 1000 system (Proxeon Biosystems). LC-MS/MS was conducted using a 120 min reversed-phase buffer gradient running at 250 nl/min (column heated to 40 °C) on a Proxeon EASY-nLC pump in-line with a hybrid LTQ-Orbitrap velos mass spectrometer (Thermo Fisher Scientific). A parent ion scan was performed in the Orbitrap, using a resolving power of 60000. Simultaneously, up to the twenty most intense peaks were selected for MS/MS (minimum ion count of 1000 for activation) using standard CID fragmentation. Fragment ions were detected in the LTQ. Dynamic exclusion was activated such that MS/MS of the same *m/z* (within a 10 ppm window, exclusion list size 500) detected three times within 45 s were excluded from analysis for 30 s.

### MS Data Processing and Protein Identification

Raw MS data were processed using MaxQuant (v1.6) with the integrated Andromeda search engine. Spectra were searched against the mouse UniProt reference proteome (UP000000589). Peptide identification was performed by matching experimental MS/MS spectra to theoretical spectra generated by in silico tryptic digestion of the FASTA database. Search parameters included trypsin specificity with up to two missed cleavages. Carbamidomethylation of cysteine was set as a fixed modification, while methionine oxidation was included as a variable modification. Minimum peptide length was set to six amino acids. Precursor and fragment mass tolerances were set to 20 ppm and 0.5 Da, respectively. The false discovery rate (FDR) was controlled at 1% at both peptide and protein levels using a target-decoy approach. The “match between runs” feature was enabled. LFQ minimum ratio count and minimum ratio count were set to 1 to improve quantification coverage.

### Protein Quantification and Statistical Analysis

A total of 249 proteins were identified. Known contaminants identified by MaxQuant were removed prior to analysis. Only proteins detected in at least 2 out of 3-4 biological replicates per condition were retained for downstream analysis. Additionally, proteins detected in more than two replicates in the GRM5^-/-^ group but absent from comparison groups were excluded to reduce bias. LFQ intensity values were log2-transformed. Zero values were treated as missing and imputed using Perseus-based normal distribution imputation. Data were then grouped by experimental condition (6-month-old wild-type males n=4, 6-month-old GRM5^-/-^ males n=3, 6-month-old wild-type females n=3, 6-month-old GRM5^-/-^ females n=3), and mean protein intensities were calculated per group. For visualization, Z-score normalization was applied to log2-transformed protein intensities across conditions to highlight relative abundance patterns independent of absolute expression levels. Z-scored data were used exclusively for clustering and heatmap visualization. After imputation, differential protein abundance was assessed at the protein level using one-way ANOVA followed by Tukey’s post-hoc test for pairwise comparisons. Multiple testing correction was applied, and p-values were used to define statistical significance. Selected biologically relevant contrasts (e.g., wild-type vs GRM5^-/-^ and sex-based comparisons) were extracted. Log2 fold changes were calculated as the difference between mean log2 LFQ intensities of the two groups. Statistical significance was defined based on adjusted p-values.

### Co-immunoprecipitation

To examine protein associations, cortical tissue was lysed in a non-denaturing buffer containing 10 mM Tris, 150 mM NaCl, 0.5 mM EDTA, and 0.5% Triton X-100, supplemented with protease inhibitor cocktail and phosphatase inhibitors (10 mM NaF and 500 μM Na₃VO₄). Lysates were gently rotated for 1 h at 4 °C and then centrifuged at 12,000 × g for 10 min at 4°C to remove insoluble material. Primary antibodies against mGluR5 (2.5 μL) or TRIM9 (2 μL) were incubated with pre-washed Protein G Mag Sepharose beads, and the mixture was rotated for 2 h at 4 °C to allow antibody binding. Pre-cleared lysate (500 μg total protein) was then added to the antibody–bead complexes and incubated overnight at 4°C with continuous rotation. Following incubation, bead-bound proteins were eluted and separated by SDS-PAGE, followed by immunoblotting to detect co-immunoprecipitated proteins, including TRIM9, PrP^C^, and mGluR5, depending on the experimental condition. In parallel, aliquots of lysates collected prior to antibody incubation were analyzed by immunoblotting to assess total levels of TRIM9, PrPᶜ, and mGluR5.

### Transfection of HEK293 cells with mGluR5 and TRIM9

Human embryonic kidney 293T (HEK293T) cells were maintained in Dulbecco’s Modified Eagle Medium (DMEM) supplemented with 10% fetal bovine serum, 100 U/mL penicillin, and 100 μg/mL streptomycin at 37°C in a humidified incubator with 5% CO₂. Cells were passaged upon reaching 85–95% confluency and used at low passage numbers. For transfection, cells were plated at ∼3 × 10⁶ cells in 100 mm dishes and incubated overnight. Transfection was performed using enhanced polyethylenimine (ePEI) at a 4:1 ratio of ePEI to DNA in Opti-MEM. Plasmids encoding FLAG-tagged mGluR5a (2 µg) and TRIM9 (2 µg) with or without 2 µg plasmid encoding G protein-coupled receptor kinase 2 (GRK2) constructs were used as indicated, and total DNA amounts were equalized using empty vector where necessary^33^. Transfection mixtures were incubated at room temperature for 20 min and added dropwise to cells. Site-directed mutagenesis was performed to substitute lysine residues at positions 690–692 within the intracellular loop 2 (IL2) region of mGluR5 (K690R, K691R, K692R; 3K→R mutant).

### Co-immunoprecipitation assay in HEK293 cells

For immunoprecipitation experiments, cells were trypsinized 18-20 h post-transfection and replated to ensure uniform conditions. Experiments were conducted 48 h post-transfection. On the day of the experiment, cells were washed with ice-cold PBS and lysed in 0.1% Triton X-100 lysis buffer (25 mM HEPES, 300 mM NaCl, 1.5 mM MgCl_2_, 0.2 mM EDTA) supplemented with protease inhibitor cocktail and phosphatase inhibitors (10 mM NaF and 500 μM Na₃VO₄). Lysates were incubated at 4 °C with rotation and clarified by centrifugation. Equal amounts of protein (500 μg) were incubated with anti-FLAG M2 affinity agarose beads at 4°C overnight with constant rotation to capture FLAG-tagged mGluR5-containing complexes. Beads were washed multiple times with lysis buffer followed by PBS to minimize non-specific binding. Bound proteins were eluted by incubation in 3× SDS loading buffer containing β-mercaptoethanol at 90 °C for 10 min. Eluted samples were resolved by SDS-PAGE and analyzed by immunoblotting using antibodies against mGluR5, TRIM9, and GRK2.

### Radioligand binding

Radioligand binding assays were performed as previously described^8,48^. Mouse cortical tissue was homogenized in ice-cold lysis buffer containing 10 mM Tris-HCl (pH 7.4) and 5 mM EDTA (pH 8.0). Crude membrane fractions were prepared by two sequential centrifugation steps at 40,000 × g for 20 min at 4 °C, with the supernatant discarded after each step and the pellet resuspended in lysis buffer. The final membrane pellet was resuspended in buffer containing 62.5 mM Tris-HCl (pH 7.4) and 1.25 mM EDTA (pH 8.0) and maintained on ice. Binding reactions were carried out in duplicate by incubating membrane preparations with 3 nM [³H]-MPEP in the presence of increasing concentrations of unlabeled MPEP or Aβ oligomers in binding assay (50 mM Tris-HCl, pH 7.4, 1 mM EDTA, pH 8.0,120 mM NaCl, 4 mM MgCl_2_, 1.5 mM CaCl_2_, and 5 mM KCl) for 90 min at room temperature. Non-specific binding was defined in the presence of 10 μM MPEP. Reactions were terminated by rapid filtration through Whatman GF/C glass fiber filters using a semi-automated harvesting system (Brandel), and retained radioactivity was measured by liquid scintillation counting (Beckman). Binding data were analyzed using GraphPad Prism software. Specific [³H]-MPEP binding was expressed as a percentage of maximal binding, and equilibrium dissociation constants (Kᵢ) for MPEP and Aβ were calculated.

### Immunoblotting

Cortical tissue was lysed in ice-cold lysis buffer containing 50 mM Tris (pH 8.0), 150 mM NaCl, 1% Triton X-100, and 0.1% sodium dodecyl sulfate (SDS), supplemented with protease inhibitor cocktail and phosphatase inhibitors (10 mM NaF and 500 μM Na₃VO₄). Lysates were clarified by centrifugation at 15,000 rpm for 10 min at 4 °C. Protein samples were mixed with lysis buffer and 5× loading buffer containing β-mercaptoethanol, then heated at 90°C for 5 min. Samples were separated by SDS-PAGE and transferred onto nitrocellulose membranes (Bio-Rad). Membranes were blocked for 1 h at room temperature in Tris-buffered saline (pH 7.6) containing 0.05% Tween-20 (TBST) and 5% nonfat dry milk. Blots were then incubated overnight at 4 °C with primary antibodies diluted 1:1000 in TBST containing 2% nonfat dry milk. Following primary antibody incubation, membranes were incubated with appropriate HRP-conjugated secondary antibodies (anti-rabbit or anti-mouse) diluted 1:7000 in TBST containing 2% non fat dry milk for 1 h at room temperature. Protein bands were visualized and quantified using Clarity Western ECL substrate (Bio-Rad, 1705061).

### Immunostaining

One hemisphere of each brain sample was fixed in 4% paraformaldehyde and then transferred to 70% ethanol for storage at 4°C. The samples were embedded in paraffin and then coronally sectioned through the hippocampus at a thickness of 5 µm. For immunofluorescence, paraffin embedded sections were deparaffinized and pre-treated using heat mediated antigen retrieval with EDTA buffer (pH 9.0). Slides were then rehydrated in TBST and blocked for 30 minutes with Rodent Block M (Biocare RBM9616). Sections were then incubated with mouse anti-Amyloid β (1:300) antibodies overnight at 4 °C. Sections were washed with TBST and then incubated with Goat anti-mouse Alexa Fluor ^TM^ 647 for 2 hours in the dark at room temperature. This was followed by incubation with a quencher (Vector TrueView Autofluorescence Quenching Kit SP-8400, Vector Labs) to decrease autofluorescence. Sections were then washed, incubated with 5 µg/ml of DAPI and coverslipped. Slides were scanned using a Leica Aperio Slide scanner (Wetzler, Germany) at 20×. For each mouse brain, one 1625*1625 µm regions of interest was analyzed per hippocampus across six brain slices to assess plaque density^49^.

### Determination of Aβ oligomer by sandwich ELISA

Levels of oligomeric Aβ were quantified using a sandwich ELISA approach as previously described^2^. Brains were dissected into right and left hemispheres, and one hemisphere was used for the measurement of oligomeric Aβ. Tissue samples were homogenized and centrifuged at 100,000 × g for 1 h at 4 °C to obtain soluble fractions enriched in Aβ oligomers. The resulting supernatant was collected and diluted 1:10 in the assay buffer provided with the kit. Quantification of oligomeric Aβ was performed using a commercially available sandwich ELISA kit (Thermo Scientific, #KHB3491) according to the manufacturer’s instructions. All samples were analyzed in triplicate. Protein concentrations were determined for each sample, and Aβ levels were normalized to total protein content.

### Statistical analysis

Statistical analyses were performed using GraphPad Prism. Data are presented as mean ± SEM. Comparisons between groups for biochemical and immunoblotting experiments were conducted using one-way or two-way analysis of variance (ANOVA), as appropriate, followed by Tukey’s post hoc test for multiple comparisons. Normality and homogeneity of variance were assessed using Kolmogorov–Smirnov and Bartlett’s tests, respectively. For mass spectrometry data, statistical analysis of differential protein abundance was performed on log2-transformed LFQ intensity values using one-way ANOVA followed by Tukey’s post hoc test with multiple testing correction applied. Details of statistical tests used for each experiment, including sample size (n) and F values, are provided in the corresponding figure legends. A value of P < 0.05 was considered statistically significant.

## Author contributions

Conceptualization: S.S.G.F. and K.S.A. Methodology: F.B. H.C. Formal analysis: F.B., R.C., Investigation: F.B., F.P., T.L.C., H.C. and R.C. Resources: M.T., S.A., and S.L.G. Data curation: F.B. Writing—original draft: F.B. Writing—review and editing: S.S.G.F. Supervision and Project administration: S.S.G.F. Funding Acquisition: S.S.G.F. and K.S.A.

## References

1. J. W. Um, et al., Metabotropic glutamate receptor 5 is a coreceptor for Alzheimer Aβ oligomer bound to cellular prion protein. Neuron 79, 887–902 (2013).

2. A. Hamilton, J. L. Esseltine, R. A. DeVries, S. P. Cregan, S. S. G. Ferguson, Metabotropic glutamate receptor 5 knockout reduces cognitive impairment and pathogenesis in a mouse model of Alzheimer’s disease. Mol. Brain 7, 40 (2014).

3. L. T. Haas, S. M. Strittmatter, Oligomers of amyloid β prevent physiological activation of the cellular prion protein–metabotropic glutamate receptor 5 complex by glutamate in Alzheimer disease. J. Biol. Chem. 291, 17112–17121 (2016).

4. L. T. Haas, et al., Metabotropic glutamate receptor 5 couples cellular prion protein to intracellular signalling in Alzheimer’s disease. Brain 139, 526–546 (2016).

5. A. Hamilton, M. Vasefi, C. Vander Tuin, R. J. McQuaid, H. Anisman, S. S. G. Ferguson, Chronic pharmacological mGluR5 inhibition prevents cognitive impairment and reduces pathogenesis in an Alzheimer disease mouse model. Cell Rep. 15, 1859–1865 (2016).

6. L. T. Haas, et al., Silent allosteric modulation of mGluR5 maintains glutamate signaling while rescuing Alzheimer’s mouse phenotypes. Cell Rep. 20, 76–88 (2017).

7. T. Bonifacino, et al., In vivo genetic ablation of metabotropic glutamate receptor type 5 slows down disease progression in the SOD1G93A mouse model of amyotrophic lateral sclerosis. Neurobiol. Dis. 129, 79–92 (2019).

8. K. S. Abd-Elrahman, et al., Aβ oligomers induce pathophysiological mGluR5 signaling in Alzheimer’s disease model mice in a sex-selective manner. Sci. Signal. 13, eabd2494 (2020).

9. K. S. Abd-Elrahman, A. Hamilton, A. Albaker, S. S. G. Ferguson, mGluR5 contribution to neuropathology in Alzheimer mice is disease stage-dependent. ACS Pharmacol. Transl. Sci. 3, 334–344 (2020).

10. K. Farmer, et al., mGluR5 allosteric modulation promotes neurorecovery in a 6-OHDA-toxicant model of Parkinson’s disease. Mol. Neurobiol. 57, 1418–1431 (2020).

11. M. Milanese, et al., Blocking glutamate mGlu5 receptors with the negative allosteric modulator CTEP improves disease course in SOD1G93A mouse model of amyotrophic lateral sclerosis. Br. J. Pharmacol. 178, 3747–3764 (2021).

12. J. A. Kos, M. Langiu, S. D. Hellyer, K. J. Gregory, Pharmacology, signaling and therapeutic potential of metabotropic glutamate receptor 5 negative allosteric modulators. ACS Pharmacol. Transl. Sci. 7, 3671–3690 (2024).

13. R. Rajmohan, P. H. Reddy, Amyloid-β and phosphorylated tau accumulations cause abnormalities at synapses of Alzheimer’s disease neurons. J. Alzheimers Dis. 57, 975–999 (2017).

14. H. Hampel, et al., The amyloid-β pathway in Alzheimer’s disease. Mol. Psychiatry 26, 5481–5503 (2021).

15. K. S. Abd-Elrahman, S. S. G. Ferguson, Noncanonical metabotropic glutamate receptor 5 signaling in Alzheimer’s disease. Annu. Rev. Pharmacol. Toxicol. 62, 235–254 (2022).

16. D. M. Walsh, et al., Naturally secreted oligomers of amyloid beta protein potently inhibit hippocampal long-term potentiation in vivo. Nature 416, 535–539 (2002).

17. P. N. Lacor, et al., Synaptic targeting by Alzheimer’s-related amyloid beta oligomers. J. Neurosci. 24, 10191–10200 (2004).

18. G. M. Shankar, et al., Amyloid-beta protein dimers isolated directly from Alzheimer’s brains impair synaptic plasticity and memory. Nat. Med. 14, 837–842 (2008).

19. J. Laurén, D. A. Gimbel, H. B. Nygaard, J. W. Gilbert, S. M. Strittmatter, Cellular prion protein mediates impairment of synaptic plasticity by amyloid-beta oligomers. Nature 457, 1128–1132 (2009).

20. M. Renner, et al., Deleterious effects of amyloid beta oligomers acting as an extracellular scaffold for mGluR5. Neuron 66, 739–754 (2010).

21. N. W. Hu, et al., mGlu5 receptors and cellular prion protein mediate amyloid-β-facilitated synaptic long-term depression in vivo. Nat. Commun. 5, 3374 (2014).

22. L. T. Haas, S. M. Strittmatter, Oligomers of amyloid β prevent physiological activation of the cellular prion protein–metabotropic glutamate receptor 5 complex by glutamate in Alzheimer disease. J. Biol. Chem. 291, 17112–17121 (2016).

23. C. Seydel, The missing sex. Nat. Biotechnol. 39, 767–769 (2021).

24. A. Waters, Society for Women’s Health Research Alzheimer’s Disease Network, M. H. Laitner, Biological sex differences in Alzheimer’s preclinical research: A call to action. Alzheimers Dement. (N. Y.) 7, e12111 (2021).

25. M. DuMont, A. Agostinis, K. Singh, E. Swan, Y. Buttle, D. Tropea, Sex representation in neurodegenerative and psychiatric disorders’ preclinical and clinical studies. Neurobiol. Dis. 184, 106214 (2023).

26. L. L. Barnes, R. S. Wilson, J. L. Bienias, J. A. Schneider, D. A. Evans, D. A. Bennett, Sex differences in the clinical manifestations of Alzheimer disease pathology. Arch. Gen. Psychiatry 62, 685–691 (2005).

27. K. S. Abd-Elrahman, A. Hamilton, M. Vasefi, S. R. Hutchinson, R. C. Russell, S. S. G. Ferguson, Autophagy is increased following both pharmacological and genetic silencing of mGluR5 signaling in mouse models of Alzheimer’s disease. Mol. Brain 11, 19 (2018).

28. K. S. Ibrahim, A. Albaker, K. S. Abd-Elrahman, S. S. G. Ferguson, Blocking estrogen receptors restores surface mGluR5 but not downstream signaling in female APP/PS1 mice. Mol. Brain 19, 68 (2026).

29. C. C. Winkle, et al., A novel netrin-1-sensitive mechanism promotes local SNARE-mediated exocytosis during axon branching. J. Cell Biol. 205, 217–232 (2014).

30. S. Menon, et al., The E3 ubiquitin ligase TRIM9 is a filopodia off switch required for netrin-dependent axon guidance. Dev. Cell 35, 698–712 (2015).

31. C. C. Winkle, et al., Trim9 deletion alters the morphogenesis of developing and adult-born hippocampal neurons and impairs spatial learning and memory. J. Neurosci. 36, 4940–4958 (2016).

32. L. E. McCormick, et al., The E3 ubiquitin ligase TRIM9 regulates synaptic function and actin dynamics in response to netrin-1. Mol. Biol. Cell 35, e23-12-0476 (2024).

33. G. K. Dhami, A. V. Babwah, R. Sterne-Marr, S. S. G. Ferguson, Phosphorylation-independent regulation of metabotropic glutamate receptor 1 signaling requires G protein-coupled receptor kinase 2 binding to the second intracellular loop. J. Biol. Chem. 280, 24420–24427 (2005).

34. Y. Chen, et al., Inhibition of mGluR5/PI3K-AKT pathway alleviates Alzheimer’s disease-like pathology through the activation of autophagy in 5XFAD mice. J. Alzheimers Dis. 91, 1197–1214 (2023).

35. Y. Qin, et al., TRIM9 short isoform preferentially promotes DNA- and RNA-virus-induced production of type I interferon by recruiting GSK3β to TBK1. Cell Res. 26, 613– 628 (2016).

36. T. Zhang, et al., G-protein-coupled receptors regulate autophagy by ZBTB16-mediated ubiquitination and proteasomal degradation of Atg14L. eLife 4, e06734 (2015).

37. K. S. Abd-Elrahman, S. S. G. Ferguson, Modulation of mTOR and CREB pathways following mGluR5 blockade contribute to improved Huntington’s pathology in zQ175 mice. Mol. Brain 12, 35 (2019).

38. G. Molinaro, et al., Female-specific dysfunction of sensory neocortical circuits in a mouse model of autism mediated by mGluR5 and estrogen receptor α. Cell Rep. 43, 114056 (2024).

39. H. Luo, et al., Amyloid-β oligomers trigger sex-dependent inhibition of GIRK channel activity in hippocampal neurons in mice. Sci. Signal. 17, eado4132 (2024).

40. S. Rubino, et al., Investigating p62 concentrations in cerebrospinal fluid of patients with dementia: A potential autophagy biomarker in vivo? Brain Sci. 12, 1414 (2022).

41. B. J. Bartlett, et al., p62, Ref(2)P and ubiquitinated proteins are conserved markers of neuronal aging, aggregate formation and progressive autophagic defects. Autophagy 7, 572–583 (2011).

42. K. S. Abd-Elrahman, A. Hamilton, S. R. Hutchinson, R. C. Russell, S. S. G. Ferguson, mGluR5 antagonism increases autophagy and prevents disease progression in the zQ175 mouse model of Huntington’s disease. Sci. Signal. 10, eaan6387 (2017).

43. M. A. Hermida, J. D. Kumar, N. R. Leslie, GSK3 and its interactions with the PI3K/AKT/mTOR signalling network. Adv. Biol. Regul. 65, 5–15 (2017).

44. J. M. Beaulieu, R. R. Gainetdinov, M. G. Caron, Akt/GSK3 signaling in the action of psychotropic drugs. Annu. Rev. Pharmacol. Toxicol. 49, 327–347 (2009).

45. J. M. Beaulieu, et al., A β-arrestin 2 signaling complex mediates lithium action on behavior. Cell 132, 125–136 (2008).

46. G. Eng, D. A. Kelver, T. P. Hedrick, G. T. Swanson, Transduction of group I mGluR-mediated synaptic plasticity by β-arrestin2 signalling. Nat. Commun. 7, 13571 (2016).

47. L. J. Stoppel, et al., β-Arrestin2 couples metabotropic glutamate receptor 5 to neuronal protein synthesis and is a potential target to treat fragile X. Cell Rep. 18, 2807–2814 (2017).

48. B. Plouffe, M. Tiberi, Functional analysis of human D1 and D5 dopaminergic G protein-coupled receptors: Lessons from mutagenesis of a conserved serine residue in the cytosolic end of transmembrane region 6. Methods Mol. Biol. 964, 141–180 (2013).

49. K. S. Abd-Elrahman, T. L. Colson, S. Sarasija, S. S. G. Ferguson, A M1 muscarinic acetylcholine receptor-specific positive allosteric modulator VU0486846 reduces neurogliosis in female Alzheimer’s mice. *Biomed*. Pharmacotherapy. 173, 116388 (2024).

